# Glymphatic Inhibition Exacerbates Tau Propagation in an Alzheimer’s Disease Model

**DOI:** 10.1101/2023.08.25.554601

**Authors:** Douglas M Lopes, Jack A Wells, Da Ma, Lauren Wallis, Daniel Park, Sophie K Llewellyn, Zeshan Ahmed, Mark F Lythgoe, Ian F Harrison

**Affiliations:** Centre for Advanced Biomedical Imaging, Department of Imaging, Division of Medicine, University College London, London, UK; Department of Internal Medicine, Section of Gerontology and Geriatric Medicine, Wake Forest University School of Medicine, Winston-Salem, North Carolina, US; Neuroscience Next Generation Therapeutics (NGTx), Eli Lilly and Company, Cambridge, Massachusetts, US

**Keywords:** Glymphatic, tau, propagation, neurodegeneration, MRI

## Abstract

**Background:** The aggregation and spread of misfolded amyloid structured proteins, such as tau and α-synuclein, are key pathological features associated with neurodegenerative disorders, including Alzheimer’s and Parkinson’s disease. These proteins possess a prion-like property, enabling their transmission from cell to cell leading to propagation throughout the central and peripheral nervous systems. While the mechanisms underlying their intracellular spread are still being elucidated, targeting the extracellular space has emerged as a potential therapeutic approach. The glymphatic system, a brain-wide pathway responsible for clearing extracellular metabolic waste from the central nervous system, has gained attention as a promising target for removing these toxic proteins.

**Methods:** In this study, we investigated the impact of long-term modulation of glymphatic function on tau aggregation and spread by chronically treating a mouse model of tau propagation with a pharmacological inhibitor of AQP4, TGN-020. Thy1-hTau.P301S mice were intracerebrally inoculated with tau into the hippocampus and overlying cortex, and subsequently treated with TGN-020 (3 doses/week, 50mg/kg TGN-020, i.p.) for 10-weeks. During this time, animal memory was studied using cognitive behavioural tasks, and structural MR images were acquired of the brain *in vivo* prior to brain extraction for immunohistochemical characterisation.

**Results:** Our findings demonstrate increased tau aggregation in the brain and transhemispheric propagation in the hippocampus and cortex following the inhibition of glymphatic clearance. Moreover, disruption of the glymphatic system aggravated recognition memory in tau inoculated mice and exacerbated regional changes in brain volume detected in the model. When initiation of drug treatment was delayed for several weeks post-inoculation, the aforementioned alterations were attenuated.

**Conclusions:** These results indicate that by modulating AQP4 function and, consequently, glymphatic clearance, it is possible to modify the propagation and pathological impact of tau in the brain, particularly during the initial stages of the disease. These findings highlight the critical role of the glymphatic system in preserving healthy brain homeostasis and offer valuable insights into the therapeutic implications of targeting this system for managing neurodegenerative diseases characterized by protein aggregation and spread.

## Background

Aggregation of misfolded amyloid structured proteins such as tau and α-synuclein can lead to the formation of amyloid fibrils and deposition of toxic insoluble cellular aggregates, which are one of the pathological hallmarks associated with neurodegenerative disorders including Alzheimer’s disease (AD) and Parkinson’s disease (1, 2). Though molecularly and structurally distinct, these amyloid structured proteins have a vicious ‘prion-like’ property in common, which gives them the unique ability to ‘seed’ and spread, allowing their transmission from cell to cell and their propagation throughout the central and peripheral nervous systems (1, 2). Although the mechanisms underlying the spread of these intracellular proteins is still being understood, mounting evidence suggests that targeting the extracellular space might represent a potential therapeutic target to manage these toxic proteins and their seeding ability in the context of neurodegenerative diseases (2).

The glymphatic system is a brain-wide pathway involved in clearance of metabolic waste from the extracellular space of the central nervous system (CNS) (3, 4), and is among potential new targets to delay or prevent clinical symptoms associated with neurodegenerative diseases. This system is modelled on the inflow of cerebrospinal fluid (CSF) along perivascular spaces surrounding arteries, which then disperses throughout the brain interstitium via exchange with interstitial fluid (ISF), resulting in efflux of fluid and interstitial solutes via veinous perivascular spaces in the brain (5).

Importantly, the glymphatic system relies on glial cells for fluid movement, and the associated interstitial solute clearance is facilitated by the water channel aquaporin-4 (AQP4), which is predominantly expressed on astrocytic endfeet ensheathing the cerebral vasculature (4, 5). Crucially, the endfeet of astroglial cells ensheath ∼99% of the cerebrovascular system (6), providing a route for the rapid movement of water between the perivascular space and the glial syncytium, and a crucial fluid flow route for solute clearance via the glymphatic pathway (7). Several studies have demonstrated that astrocytic AQP4 is essential for glymphatic function, with AQP4 knockout animal models exhibiting severe disruption of CSF-ISF exchange and solute clearance (3, 7–12). Further, it has been demonstrated that the glymphatic system is involved in the clearance of amyloid-β (Aβ), the main component of the amyloid plaques found in the brains of people with AD (3, 13), and mounting evidence also links the glymphatic system with the clearance of the intracellular AD hallmark protein tau (8, 11, 14).

Many studies demonstrate that tau can be secreted to the extracellular space and taken up by both neurons and glial cells, where it plays an important role in the development of tauopathies (2, 15–17), indicating a potential role the glymphatic system might play in clearing away extracellular tau prone to pathogenesis (14). Indeed, strong evidence suggests that interstitial tau is cleared via the same perivascular pathways as Aβ, along the glymphatic network (11). Further, it has been shown that p-tau accumulation in the mouse brain is inversely correlated with the expression of AQP4, with AQP4 knockouts exhibiting an increased level of intracellular p-tau, exacerbated neuronal degeneration and cognitive dysfunction (8). In addition, we recently demonstrated that AQP4 polarisation and glymphatic function were reduced in the brain of a transgenic mouse model of tauopathy, and that acute pharmacological blockade of AQP4 resulted in disrupted tau clearance from the brain (14). These data support the notion of a strong relationship between impaired glymphatic function and tau deposition in the rodent brain, indicating that inadequate clearance of interstitial tau, prone to prion-like propagation and spread, likely contributes to tau aggregation (2). Furthermore, glymphatic function has been shown to become impaired with age, the greatest risk factor for dementia, with widespread loss of AQP4 polarisation along penetrating arteries (18, 19).

To explore the link between glymphatic function and tau propagation, we set out to determine whether long term modulation of glymphatic function would affect tau aggregation and spread. Using an established model of tau propagation (Thy1-hTau.P301S mice intracerebrally inoculated with tau (20)), we chronically treated mice with a pharmacological AQP4 inhibitor, TGN-020, and evaluated the extent of tau pathology and spread, together with the resultant effects on regional brain atrophy, and changes in behaviour. We found that mice chronically treated with the1AQP4 inhibitor showed a greater extent of tau aggregation and propagation in the brain following chronic glymphatic impairment. Moreover, these animals exhibited amplified cognitive impairments during memory tasks, along with intensified brain atrophy, indicating that the modulation of AQP4 function and, consequently, the glymphatic system can significantly influence the propagation of tau throughout the brain. Additionally, these findings imply that by targeting the function of AQP4 and the glymphatic system, it is possible to alter the extent to which tau is capable of propagating throughout the brain, thereby influencing the development of pathology and neurodegeneration. By extension, these data highlight the vital role the glymphatic system plays in healthy brain homeostasis.

## Methods

### Animals

Generation of homozygous Thy1-hTau.P301S transgenic mice (line 2541, henceforth referred to as P301S mice) has been reported previously (21). Mice were bred on a mixed C57BL/6JxCBA/ca background, and breeders transferred from the MRC Laboratory of Molecular Biology (Cambridge, UK) for colony establishment at UCL (London, UK). For the non-transgenic (wildtype) experiments, C57BL/6J mice were acquired from Charles River, UK. Animals were housed on a 12-hr light/dark cycle in groups of two to five in individually ventilated cages with *ad libitum* access to food and water and provided with environmental enrichment (cardboard houses and tubes, paper nesting). Equal numbers of both male and female animals were used in experimental groups. All animal work was performed in accordance with the UK’s Animals (Scientific Procedures) Act of 1986 and approved by UCL’s internal Animal Welfare and Ethical Review Board.

### Cisterna Magna Infusion for Assessment of Glymphatic Function

Glymphatic function was assessed in the mouse brain by infusion of a fluorescent tracer via the cisterna magna, followed by whole brain and brain slices imaging for quantification of CSF-brain influx, similar to that previously published (3). Briefly, mice were anesthetized with 2% isoflurane in O_2_ at a delivery rate of 1 l/min and positioned in a stereotaxic frame with the head flexed to ∼50°. A midline incision was made at a midpoint between the skull base and the occipital margin to the first vertebrae. The underlying muscles were parted to expose the atlanto-occipital membrane and dura mater overlaying the cisterna magna, and a durotomy was performed using a 23-gauge needle. An intrathecal catheter (35–40 mm port length, 80–90 mm intravascular tippet length, Braintree Scientific, US) extended with polyethylene tubing (0.4 mm × 0.8 mm, Portex™, Smiths Medical, UK) and attached to a 100 µl glass Hamilton syringe driven by a microinfusion pump (sp210iw syringe pump, World Precision Instruments, US) was filled with low molecular weight fluorescent tracer, Texas Red™-conjugated dextran (TxR-d3, 3 kDa; ThermoFisher Scientific, UK) at 0.05% in filtered artificial CSF (Tocris, UK), advanced 1 mm into the cisternal space, and sealed and anchored in place with cyanoacrylate. Once dry, TxR-d3 was infused via the implanted cannula (2 µl/min over 5 min). 30 min after the start of the infusion, mice were killed by overdose with sodium pentobarbital (10 ml/kg, given intraperitoneally) and the brain was quickly removed.

Whole brain fluorescence was imaged immediately in excised brains (Ex. 570 nm, Em. 600 nm) using an IVIS Spectrum imaging system (PerkinElmer, UK). Brains were then drop-fixed in 4 % paraformaldehyde (PFA) for 48 hrs, prior to cryoprotection (30 % sucrose in PBS for 72 hrs), freeze embedding in OCT, and sectioning (100 µm sagittal sections) onto SuperFrost Plus™ microscope slides (VWR, UK) using a cryostat (CM3050 S, Leica, UK). Slices were mounted using Fluoromount-G^TM^ with DAPI (Invitrogen, UK), and single plane images were captured on a Leica microscope (Leica DMi8) using a 10× 1.3 NA objective with Texas Red and DAPI filter sets. Whole brain fluorescence in three aspects (dorsal, ventral and lateral) was measured using the Living Image Analysis Software (PerkinElmer, UK), and area coverage of Texas Red fluorescence in brain slice images was measured using ImageJ (v1.51j8).

### Brain Extracts

Brain extracts for subsequent intracerebral infusion were prepared from end-stage (5-5.5 months-of-age) P301S and age-matched wildtype mice, as previously described (20). Briefly, mice were killed by overdose with sodium pentobarbital (10 ml/kg, given intraperitoneally), brainstems were rapidly dissected on wet ice and tissue extracts snap frozen on dry ice for storage at -80 °C. Frozen brainstems from five mice were combined, weighed, and homogenised in 10% (w/v) sterile phosphate-buffered saline (PBS) (containing protease inhibitor cocktail, phosphatase inhibitor cocktails I and II (Sigma, UK), at a final dilution of 1:100, and 1 mM phenylmethylsulphonyl fluoride) by sonication (Fisherbrand™ Model Sonic Dismembrator, Fisher Scientific, UK). The resultant brain extract homogenates (henceforth referred to as either ^P301S^BE or ^WT^BE, originated from P301S or wildtype animals, respectively) were centrifuged at 3,000 g at 4 °C for 5 min, and the supernatants stored at -80 °C until characterisation and subsequent intracerebral infusion.

Total tau content was quantified in brain extracts by ELISA (Human Tau (total) ELISA kit (#KHB0041, Invitrogen, UK), see supplementary methods) (31.2(±2.8) µg/ml in ^P301S^BE, 0.1(±2.4) µg/ml in ^WT^BE), and dot blots performed (see supplementary methods) to confirm immunoreactivity of ^P301S^BE (and not ^WT^BE) with MC1 (abnormal AD tau conformation) and TG5 (total tau) antibodies (both a kind gift from the late Prof Peter Davies), and with the 4R tau specific RD4 (1E/A6) antibody (a kind gift from Prof Rohan de Silva (22)) (**Suppl. Fig. 1**). Additionally, presence of amyloid fibrils in ^P301S^BE was verified by its fluorescence upon incubation with Thioflavin T (see supplementary methods) (signal intensity (a.u.) 1.8(±0.1) for ^P301S^BE, 0.9(±0.1) for ^WT^BE). Prior to intracerebral infusion, ^P301S^BE was adjusted to 20 μg/ml total tau (based on ELISA determined concentration) with homogenization buffer (^WT^BE diluted by the same factor), briefly sonicated, and kept on ice until infusion.

### Intracerebral Infusion of Brain Extracts

Mice (2 months-of-age) were anesthetized with 2% isoflurane in O _2_ at a delivery rate of 1 l/min and positioned in a stereotaxic frame in the horizontal skull position. A midline incision was made on the top of the head to expose the skull and a burr hole made with a microdrill above the injection location (+2.5 mm anteroposterior, and +2 mm mediolateral to bregma). 2.5 µl/site of either ^P301S^BE or ^WT^BE was infused at a rate of 0.5 µl/min into the hippocampus and overlying cortex (-1.8 and -0.8 mm ventrodorsal to the brain surface respectively) using a 5 µl Hamilton syringe. After each infusion, the needle was left*in situ* for 5 min to prevent injectate reflux. The scalp was then sutured closed, and the animal left to recover in a heated recovery chamber until it had regained consciousness. It was then returned to its home cage, and its weight monitored daily for 7 days post-surgery to ensure complete recovery.

Due to their endogenous expression of mutant human tau, ‘seeding’ of young P301S mice as above leads to robust and rapid development of tau pathology within days of inoculation, free from the confounds of regional transgene driven deposition: mild/early pathology is observable 2 weeks to 1 month following seeding, and severe/late tau pathology is evident ∼2 months (20). Of note, this phenomenon is specific to transgenic P301S mice, as when wildtype mice were injected with ^P301S^BE and left to age for 10-weeks (either treated with vehicle or TGN-020, see below), brain tau pathology remained barely detectable by histology ( **Suppl. Fig. 2**). AQP4 expression (see supplementary methods) also appears to be unaffected by disease onset in ^P301S^BE infused transgenic mice, supporting its targeting in this model for mechanistic study, as we have done here ( **Suppl. Fig. 3**).

### Drug Treatments

Pharmacological inhibition of AQP4 was achieved using TGN-020 (N-1,3,4-thiadiazol-2-yl-3-pyridinecarboxamide, Tocris Bioscience, UK). Unless otherwise stated, mice were treated intraperitoneally with either TGN-020 (50 mg/kg in 20 ml/kg 0.9% NaCl (saline)) or vehicle (20 ml/kg saline). For chronic treatment studies, animals received injections of either TGN-020 or vehicle, 3 times per week during a 10-week period, starting ∼3 days post-surgery, unless otherwise stated. In control (wildtype) animals, this chronic treatment regimen had no negative or confounding effects on the animals’ health, body weight ( **Suppl. Fig. 4A**), cognitive performance (**Suppl. Fig. 4B**), or locomotor ability (**Suppl. Fig. 4C**) throughout treatment. And neither expression nor polarisation of AQP4 was affected by TGN-020 treatment in control animals ( **Suppl. Fig. 5**)

### Immunohistochemistry

Mice were injected with an overdose of sodium pentobarbital (10 ml/kg, given intraperitoneally) and transcardially perfused with approximately 10 ml PBS immediately followed by the same volume of 4 % PFA in PBS. Brains were extracted from the skull and drop-fixed in 4 % PFA for 24 hrs at 4 °C. Brains were then cryopreserved (30 % sucrose in PBS for 72 hrs at 4 °C), freeze embedded in OCT, and serial coronal sections collected (50 µm thick) onto SuperFrost Plus™ microscope slides (VWR, UK) using a cryostat (CM3050 S, Leica, UK). Slides were stored at -20 °C until further processing. On the day of staining, sections were brought to room temperature (RT) before blocking in 10 % donkey serum (in 0.02 % Triton X-100 PBS) for ∼4 hrs at RT. Sections were then incubated with primary antibody (mouse anti-phospho-Tau (AT8, 1:500, Invitrogen, UK), rabbit anti-NeuN (D3S3I, 1:600, Cell Signalling, UK) or rabbit anti-AQP4 (1:500, Millipore, UK)), overnight at RT in blocking buffer. Slides were washed three times and incubated with secondary antibody (anti-mouse or anti-rabbit (depending on primary antibody host) conjugated to Alexa Fluor™ 568 (1:800, donkey, Invitrogen, UK)) for ∼4 hrs at RT in the dark. Slides were then washed again three times and mounted under coverslips using Fluoromount-G ^TM^ with DAPI (Invitrogen, UK). Slides were kept in the dark until imaging.

Single plane, tiled images were captured on a fluorescence microscope (Leica DMi8) using a 10× 1.3 NA objective. For analysis, using ImageJ (v1.51j8), a region-of-interest (ROI) was drawn on images encompassing the target region (e.g. hippocampus) in at least 6 sections from each brain, with the experimenter blind to animal experimental/treatment group. Measures of signal intensity and % area coverage (after thresholding for immunoreactivity) were extracted from each and averaged across slides analysed from each brain. The interhemispheric area ratio (contralateral % area coverage divided by the ipsilateral % area coverage) was additionally calculated, as an indicator of transhemispheric propagation.

### Behavioural Experiments

#### Open Field

The apparatus consisted of a 45 x 45 x 45 cm white Perspex box open at the top. At the start of the test the mouse was placed in the centre of the field, and allowed to explore the apparatus for 5 min. The open field was thoroughly cleaned with 70% ethanol in between trials. Each mouse was recorded using a digital video camera (Konig, CASC300) placed approximately 50 cm above the apparatus. Animals’ paths were tracked using AnimalTracker ( http://animaltracker.elte.hu/), which automatically determined the distance travelled and time spent in the central and peripheral areas of the field. Path tracking analysis was carried out with the experimenter blinded to experimental/treatment group.

#### Novel Object Recognition (NOR)

NOR testing took place in the open field arena (described above) after the animals had been acclimatised to the field. At the beginning of the experiments two identical objects were placed inside, in a diagonal position, equidistant to the walls. Mice were allowed to freely explore and habituate to the two identical objects for 5 min and then removed from the field. Following a retention interval (24 hrs), mice were placed back into the arena with the objects in the same locations but with one of the familiar objects replaced with a novel object, the latter being different in shape, size, colour and texture. Mice were then allowed to freely explore both objects for 5 min. Animals were recorded using a digital camera positioned above the arena throughout the experiment, as above. Objects were thoroughly cleaned with 70% ethanol before and in between each trial. A mouse was considered to be exploring the object when its head was facing the object at a distance of approximately 2 cm or less or when part of its body except the tail was touching the object (in the event of an animal climbing the object, the time was not counted as exploration time). Scoring following the retention interval was carried out manually with the experimenter blinded to experimental/treatment group. Preference index (PI) was calculated based on the time the animal spent exploring the novel object divided by the total time exploring both objects:

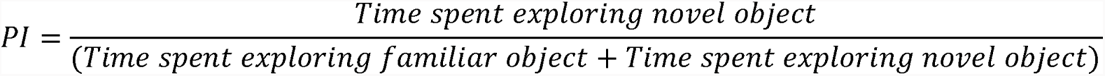

#### Hindlimb Clasping Scoring

Assessment of corticospinal function with limb clasping was performed based on images captured using a portable hand-held device (smart phone). Each animal was removed from its home cage, gently suspend by the tail just above a flat surface (metal grid) and an image was acquired whilst the animal’s hind paws were suspended. Images were scored following the pre-defined criteria with the experimenter blinded to experimental/treatment group (23):

1. No limb clasping. Normal escape extension. One hind limb exhibits incomplete splay and loss of mobility. Toes exhibit normal splay.
2. Both hind limbs exhibit incomplete splay and loss of mobility. Toes exhibit normal splay.
3. Both hind limbs exhibit clasping with curled toes and immobility.
4. Forelimbs and hind limbs exhibit clasping and are crossed, curled toes and immobility.

### MRI Scanning

At the end of drug treatment studies (10-weeks post-inoculation), structural 3D MR images of mouse brains were acquired. For this, mice were anesthetized with 2% isoflurane in a mixture of air (0.4 l/min) and O_2_ (0.1 l/min), and positioned into an MR-compatible stereotaxic frame, with 1.5% isoflurane delivered (in the same gas mixture) via a nose cone. A horizontal-bore 9.4T Bruker preclinical system (BioSpec 94/20 USR, Bruker, Germany) was used for acquisition of 3D structural T2-weighted MR images of the entirety of the mouse brain, using a volume coil and 4-channel surface coil for RF transmission and signal detection, respectively. A fast spin echo sequence with the following parameters: FOV = 16 x 16 x 16mm, matrix = 128 x 128 x 128, repetition time = 700ms, echo time = 40ms, 2 averages.

For analysis, a multi-atlas brain structure parcellation framework was used (24) to extract the whole brain structural volumes. The raw brain MR images went through the pipeline of reorientation, intensity non-uniformity correction through the N4 bias correction algorithm, skull stripping, and whole-brain segmentation. The template images of an *in vivo* 3D digital atlas of the adult C57BL/6J mouse brain (25) were first registered affinely to the original MR image data using a block-matching algorithm (26). The brain masks in the atlas were then transformed and resampled to create a consensus brain mask. A further non-rigid registration based on fast free-form deformation was then performed to correct any remaining local misalignment of the affinely registered atlas to the brain volumes (27). The brain structure labels from the atlas were then transformed and resampled to the individual image space transformation and fused using the STEPS label fusion algorithm (28) to create the final parcellation. Hemispheric volumes were extracted for interrogation of interhemispheric differences, calculating the degree of interhemispheric volume ratio (ipsilateral/contralateral volume) change from 1 (hemispheric equality), and comparing this across groups/regions to determine if the degree of interhemispheric volume difference was altered with drug treatment in any brain region.

### Statistical Analysis

All data is presented as group means ± SEM, with individual animal datapoints shown where possible. Unpaired t-tests were used for comparisons made between two groups. For comparison of three or more groups, either 1-way, 2-way or 3-way ANOVA were used, depending on the dataset (ANOVA type denoted on individual graphs), with either Tukey’s multiple comparisons tests for individual comparisons, or Sidak’s multiple comparisons tests for grouped data. Differences in MRI data were analysed by multiple t-tests, correcting for multiple comparisons using the Holm-Sidak method, plotting -log(Adjusted P Value) against the magnitude of change of differences in volcano plots. For XY plots, the lines of best fit were smoothed using 2 neighbour datapoints. All graphs were constructed, and statistical comparisons made using GraphPad Prism (v8.1.2).

## Results

### Chronic Pharmacological Blockade of AQP4 Results in Sustained Glymphatic Inhibition

Studies have demonstrated that knockout model systems for AQP4 gene deletion reproducibly lead to significant decreases in glymphatic function and protein clearance (3, 7, 29). In addition, we have previously demonstrated that acute selective pharmacological blockage of AQP4 channels, using TGN-020 (30–32) leads to a >80% reduction in CSF-ISF exchange across different regions of the brain, and a corresponding decrease in solute clearance (14). Our initial objective was therefore to investigate whether TGN-020 would have a similar effect when administered chronically. Adult wildtype mice (2 months-of-age) were treated with either TGN-020 or vehicle over a period of 3-weeks, with 3 doses/week (50mg/kg TGN-020 in 20mg/kg saline or 20mg/ml saline alone, i.p.); concentration and dose chosen based on our drug efficacy findings in relation to brain tau clearance (see supplementary methods, **Suppl. Fig. 6**). Severe disruption of CSF-ISF exchange was observed in animals chronically treated with TGN-020 following cisterna magna infusion with TxR-d3, with only half of the fluorescent signal detected compared to vehicle-treated animals (p<0.001, **Fig. 1A, B**).

**Figure 1.**
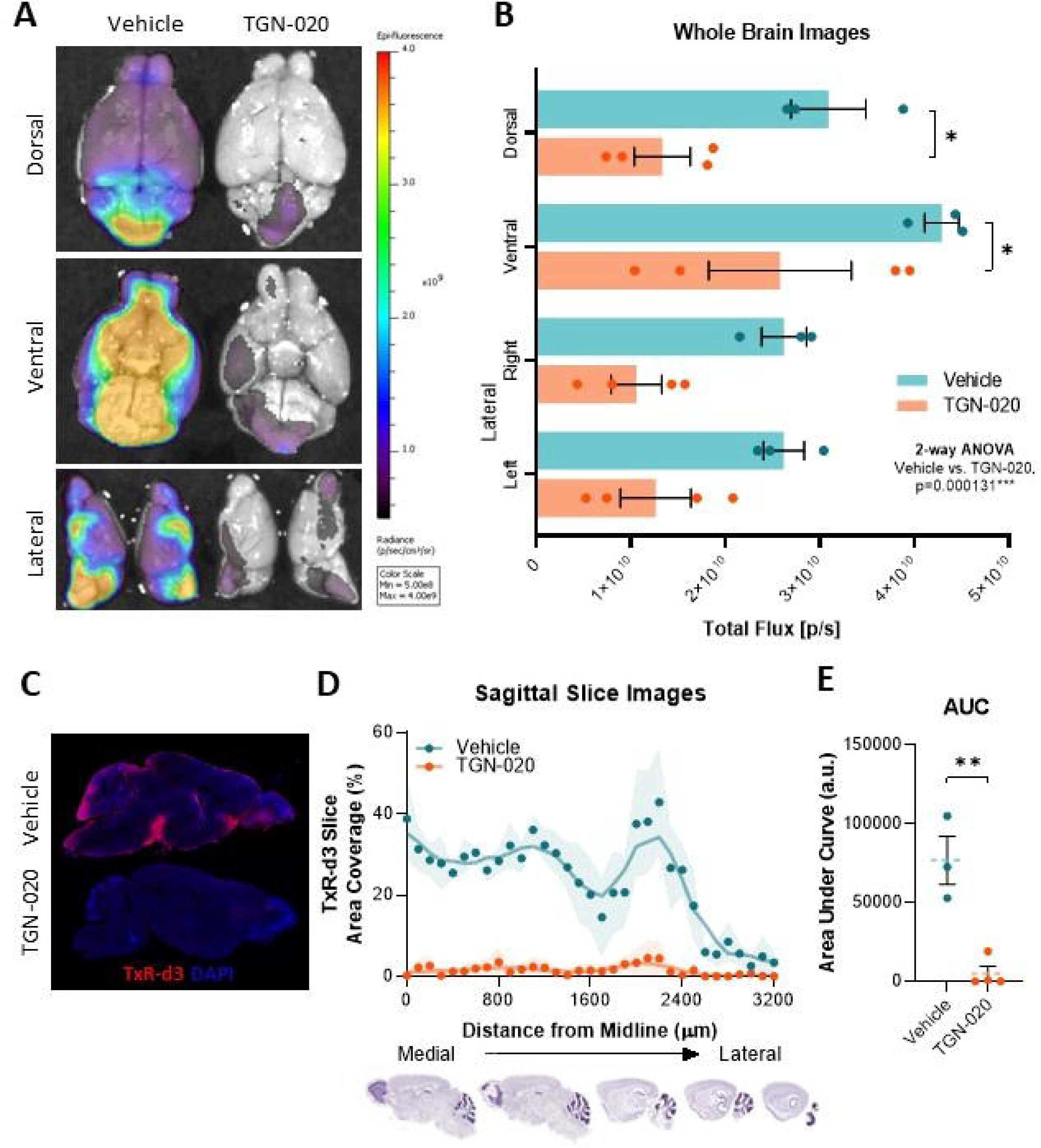
Chronic Pharmacological Blockade of AQP4 Results in Sustained Glymphatic Inhibition. **(A)** Representative example of whole brain images with psuedocoloured fluorescence overlaid, illustrating the dramatic reduction of brain surface coverage of fluorescence tracer (TxR-d3, 3kDa) following chronic treatment with TGN-020 (50 mg/kg) compared to vehicle. Quantification of this signal (**B**) shows that in all three orientations, TGN-020 treated mice exhibit a reduction in tracer coverage. Sectioning of brains and imaging by fluorescence microscopy demonstrated a similar difference; (**C**) representative examples of a brain section (∼1 mm lateral to the midline) of a vehicle treated and a TGN-020 treated mouse brain, showing clear surface penetration of TxR-d3 in the vehicle treated but not TGN-020 treated animal. ( **D**) Quantification of the % area of each section covered by the tracer demonstrates this difference more clearly (shaded area indicates SEM of animals examined). (**E**) Area under curve of data shown in ( **D**). *=p<0.05, **=p<0.01, n=3-4 per group.

Further, when sectioned, a dramatic reduction in the percentage area of brain tissue covered by the CSF tracer was observed in drug treated mice ( **Fig. 1C, D**): area under the curve of distance vs. signal plots indicated a 93(±13) % decrease in tracer influx ( **Fig. 1E**), with regions closest to the medial part of the brain (midline) being the most affected (**Fig. 1D**). These results demonstrate that chronic pharmacological blockade of AQP4 via TGN-020 treatment of adult mice significantly reduces CSF-ISF exchange and supresses brain-wide glymphatic function.

### Chronic Glymphatic Inhibition Exacerbates Tau Propagation in a Mouse Model

Recent studies suggest a relationship between tau accumulation, AQP4 polarisation and distribution, and glymphatic function (11, 14), highlighting the possible impact the latter can have on propagation of tau directly. Given this evidence, we set out to investigate the consequences of chronic glymphatic suppression, via pharmacological AQP4 blockade, on tau propagation, in an *in vivo* model setting. For this, we induced tau propagation in P301S mice, as has been previously described (20), and chronically treated animals with the AQP4 inhibitor, TGN-020, as above (3 doses/week, 50mg/kg TGN-020 in 20mg/kg saline or 20mg/ml saline alone, i.p.), for 10 weeks, to determine the resultant effects on the propagation of tau in the brain.

A clear trend of increasing intensity of tau (AT8 immunopositive) pathology was observed in the molecular layer of the hippocampus upon inoculation with tau ( ^P301S^BE, containing pathogenic tau) and with drug treatment (**Fig. 2A, B**); with TGN-020 treated ^P301S^BE injected mice exhibiting a significantly higher tau signal than ^WT^BE (not containing pathogenic tau) injected controls (p<0.05, **Fig. 2B**). ^P301S^BE injected vehicle treated mice exhibited a significant interhemispheric difference in tau immunopositive area (ipsi>contralateral, p<0.05, **Fig. 2C**) which was not seen in TGN-020 treated mice. As such, in drug treated mice, tau propagation to the contralateral hippocampus was considerably higher (relative to the ipsilateral side, p<0.01 compared to vehicle treated ^P301S^BE injected animals, **Fig. 2D**).

**Figure 2.**
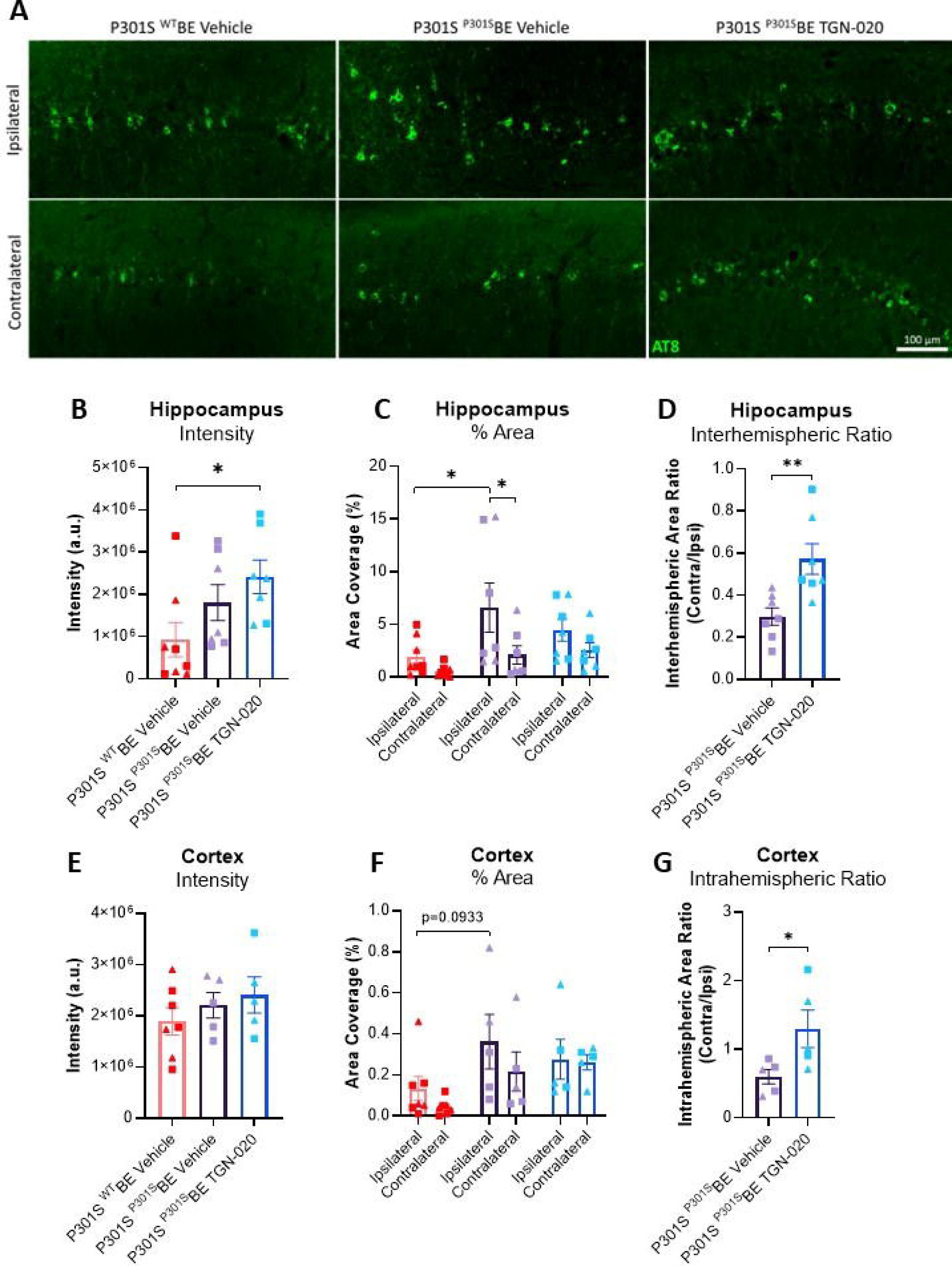
Chronic Glymphatic Inhibition Exacerbates Tau Propagation in a Mouse Model. (**A**) Representative examples of AT8 immunoreactivity in the hippocampus of treated mice groups. (**B**) Quantification of AT8 immunoreactive intensity in the hippocampus indicates exacerbated tau pathology in TGN-020 treated ^P301S^BE injected transgenic mice. When subdivided into hemispheres (**C**), ^P301S^BE infused vehicle treated mice exhibited a significant difference in immunopositive staining (% area) between hemispheres of the hippocampus, which was not evident in TGN-020 treated mice. This increase in contralateral tau deposition in TGN-020 treated mice was made more clear when calculating the interhemispheric area ratio in tau injected mice ( **D**); TGN-020 treated animals displaying far greater contralateral tau positivity in relation to the injected hemisphere. Similar, albeit more subtle differences were observed in the cortex, i.e. the other region of the brain to receive tau inoculation. A subtle trend of increase from ^WT^BE injected, to ^P301S^BE injected, to ^P301S^BE injected TGN-020 treated mice was observed in cortical AT8 intensity ( **E**). When subdivided into hemispheres, no significant differences between groups/hemispheres was observed ( **F**). When calculating the interhemispheric area ratio in tau injected mice however ( **G**), a significance increase was observed between vehicle and TGN-020 treated ^P301S^BE injected cohorts, indicative of exacerbated interhemispheric propagation of tau with drug treatment. ▴=females, ▪=males, *=p<0.05, **=p<0.01, n=5-8 per group.

A similar trend towards increasing tau signal and area coverage with both inoculation with tau (^P301S^BE) and with drug treatment was observed in the cortex (encompassing the somatosensory, visual and retrosplenial cortices, **Fig. 2E, F**). Likewise, greater propagation to the contralateral cortex was observed in TGN-020 treated mice (relative to the ipsilateral side, p<0.05 compared to vehicle treated ^P301S^BE injected animals, **Fig. 2G**).

Together, these results show that chronic suppression of glymphatic function through AQP4 inhibition in a mouse model of tau propagation, exacerbates pathological spread of tau in the brain.

### Chronic Glymphatic Inhibition Exacerbates Memory Impairment in a Mouse Model of Tau Propagation

Given that chronic pharmacological suppression of AQP4 *in vivo* leads to glymphatic impairment and increased propagation of tau, we next tested whether this led to any change in the behaviour of the animals. After only 4 weeks of TGN-020 treatment, we observed a marked decline of preference index in ^P301S^BE injected mice compared to baseline in a Novel Object Recognition (NOR) paradigm (p<0.001, **Fig. 3A**); a behavioural task dependent on the hippocampal formation and its cortical connections (33–35). This substantial decline in object preference was significantly greater than that observed in ^WT^BE injected mice (p<0.05, **Fig. 3B**), suggesting that the exacerbated effect of TGN-020 treatment on tau burden and spread has a negative impact on function of recognition memory in these animals.

**Figure 3.**
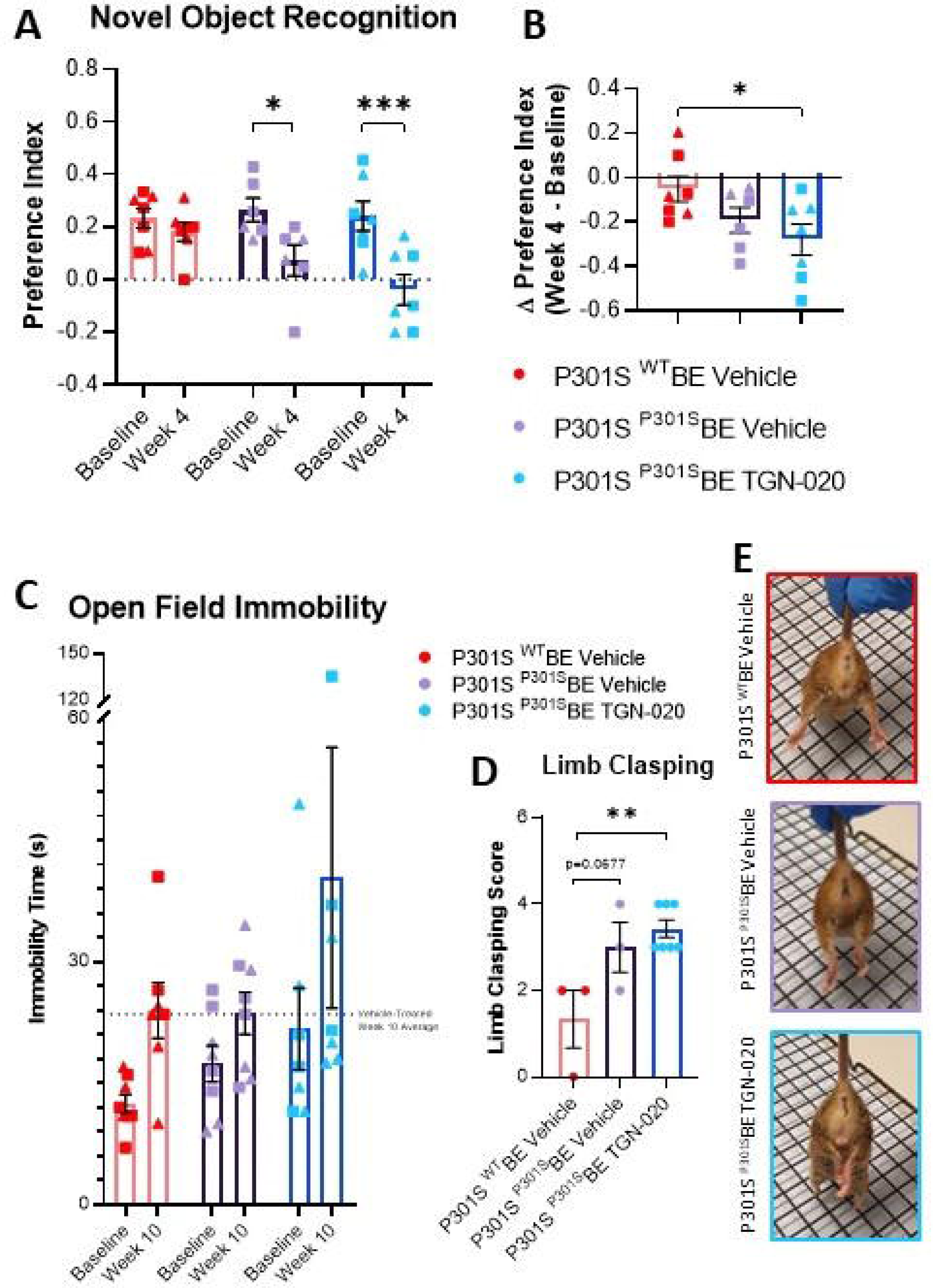
Chronic Glymphatic Inhibition Exacerbates Long-Term Memory Impairment in a Mouse Model of Tau Propagation. Following 4 weeks of TGN-020 treatment, ^P301S^BE injected mice do not exhibit any strong preference towards the novel unfamiliar object in the 24 hr delayed trial of the Novel Object Recognition test (**A**). Upon calculating the difference in preference index between week 4 and baseline (**B**), statistically significance was observed between^WT^BE injected vehicle treated and ^P301S^BE injected TGN-020 treated mice. (**C**) TGN-020 treated animals on average appear to spend more time immobile in the open field task, with almost half of the drug treated animals spending extended periods of the time immobile. Limb clasping scores in a subset of the animals ( **D**) indicate that TGN-020 treated ^P301S^BE injected mice exhibit significantly greater hind-limb clasping compared to ^WT^BE injected mice. (**E**) Representative scored example images of hind-limb clasping behaviour exhibited by animals. ▴=females, ▪=males, *=p<0.05, **=p<0.01, n=7-8 per group.

In an open field arena 10 weeks post-inoculation, TGN-020 treated mice were on average less mobile in comparison to the other groups, with almost half of the drug treated mice remaining immobile for extended periods during the task (**Fig. 3C**), despite no significant difference in total path length relative to the other groups ( **Suppl. Fig. 7A**). Further, at this same timepoint, TGN-020 treated mice exhibited the highest overall limb clasping score amongst the groups (p<0.01 compared to ^WT^BE injected animals, **Fig. 3D, E**), indicative of a substantial decline in corticospinal function in these animals. We also attempted to perform the NOR task at this 10-week timepoint following drug treatment. However, we noticed that animals treated with TGN-020 spent insufficient time exploring either of the objects (**Suppl. Fig. 7B**), and as such this timepoint could not be reliably considered, as lack of interaction and interest with the objects resulted in confounding of the test. This behaviour could be attributed to animals’ immobility and inactivity, as evident from the above datasets (**Fig. 3C-E**).

### Chronic Glymphatic Inhibition Appears to Exacerbate Volumetric Changes Caused by Tau Propagation, in Extra-Hippocampal/Cortical Regions

It is proposed that tau propagation is related to synaptic connectivity rather that spatial proximity, and indeed, it has been shown that in the mouse model used in this study, tau pathology spreads to synaptically-connected extra-hippocampal/cortical areas (20). In order to assess the extent of tau induced volume changes outside the hippocampus, we acquired structural magnetic resonance images of the whole brain and automatically parcellated them to extract volumes of gross neuroanatomical structures. For each brain region in each group, interhemispheric volume differences were examined by calculating the interhemispheric volume ratio (ipsi/contralateral volume), allowing appreciation of tau/drug induced changes both between hemispheres of a region of the brain and between brain regions.

Exacerbated interhemispheric volume differences were observed in TGN-020 treated ^P301S^BE injected mice, as is evident from volcano plotting of the differences in interhemispheric volume ratio ( **Fig. 4**). Significant ^P301S^BE injected TGN-020 treated mice datapoints exhibit both an upper-left (accentuated ipsi<contralateral difference) and upper-right (accentuated ipsi>contralateral difference) shift away from the centre of the plot compared to vehicle treated ^P301S^BE injected group points (**Fig. 4**). Not only are the degrees of significance of hemispheric differences exaggerated in TGN-020 treated mice compared to vehicle, but the magnitudes of effect also appear greater upon treatment with the drug. For example, in TGN-020 treated compared to vehicle treated ^P301S^BE injected mice, we observed a 49% and 29% exacerbation of ipsilateral atrophy in the striatum and amygdala respectively (**Suppl. Fig. 8**). Likewise, the hemispheric differences in anterior commissure and midbrain volumes (ipsi>contralateral) were magnified by 36% and 11% respectively in drug treated mice compared to vehicle (**Suppl. Fig. 8**). These data suggest that TGN-020 aggravates tau induced volume changes outside the site of initial tau seeding, indicating intensified propagation of tauopathy as a result of glymphatic inhibition in the brain.

**Figure 4.**
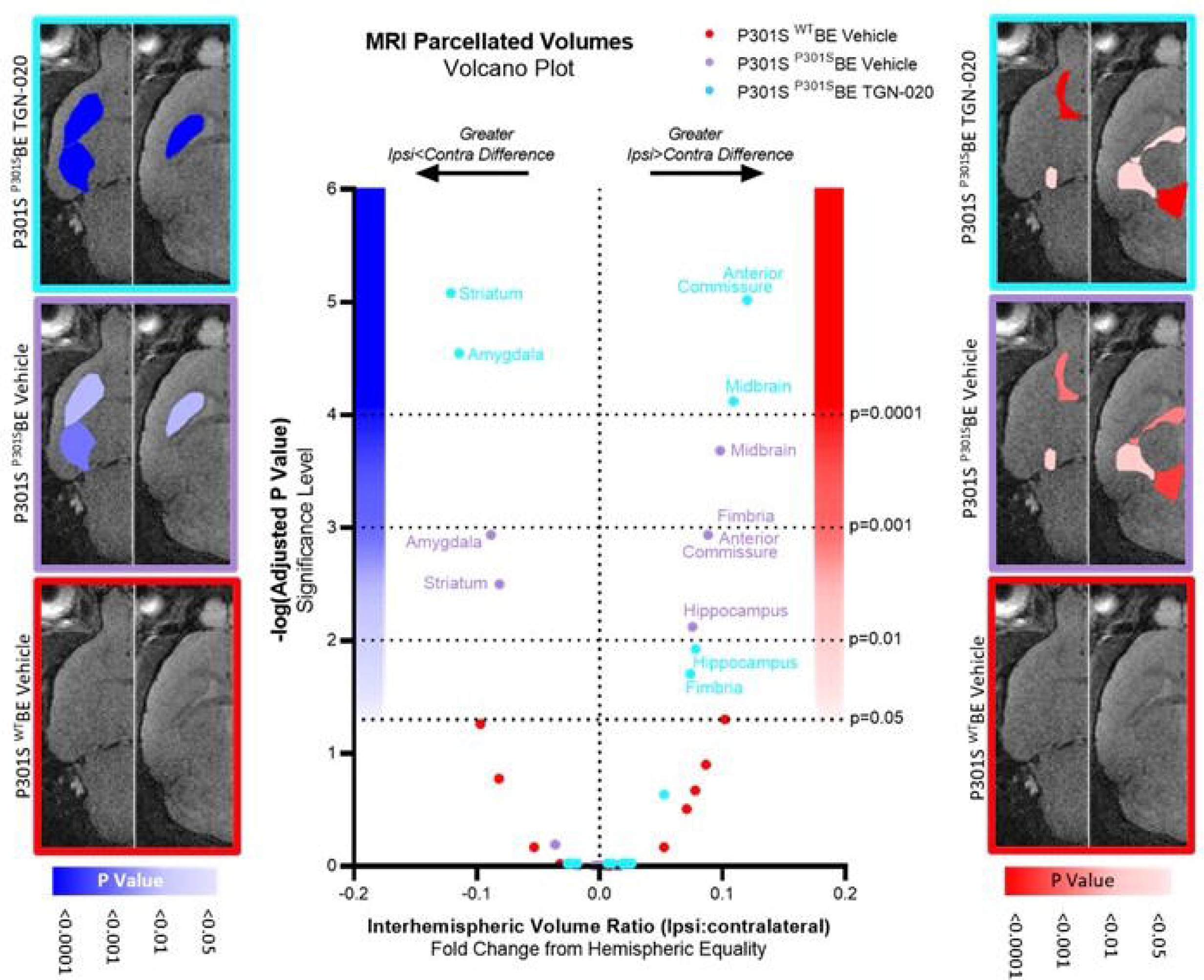
Chronic Glymphatic Inhibition Appears to Exacerbate Volumetric Changes Caused by Tau Propagation, in Extra-Hippocampal/Cortical Regions. Volcano plot of comparisons between groupwise regional interhemispheric volume ratios and hemispheric equality, indicating both an upper-left (accentuated ispi<contralateral difference) and upper-right (accentuated ispi>contralateral difference) shift away from the origin (bottom centre) of the plot in P301S ^P301S^BE TGN-020 treated group datapoints compared to vehicle treated controls. For anatomical reference, horizontal slice images are shown (left, -4 from Bregma; right, 2mm from Bregma) with regions colour scaled based on their significant interhemispheric differences (ipsi<contralateral in blue (left), ipsi>contralateral in red (right)) in each of the treatment groups.

### Delayed TGN-020 Treatment Impacts Its Effect on Tau Propagation

The above data suggests that chronic suppression of glymphatic function leads to a clear increase in the propagation of tau, resulting in discernible cerebral atrophic changes and cognitive behavioural outcomes in a mouse model of tau propagation. To investigate this phenomenon further and to gain insights into the role of the glymphatic system in disease progression, we conducted additional animal experiments in which we initiated drug treatment at either 2-weeks (early/mild stages of tauopathy) or 4-weeks (moderate stages of tauopathy) after tau inoculation (20) (**Fig. 5A**).

**Figure 5.**
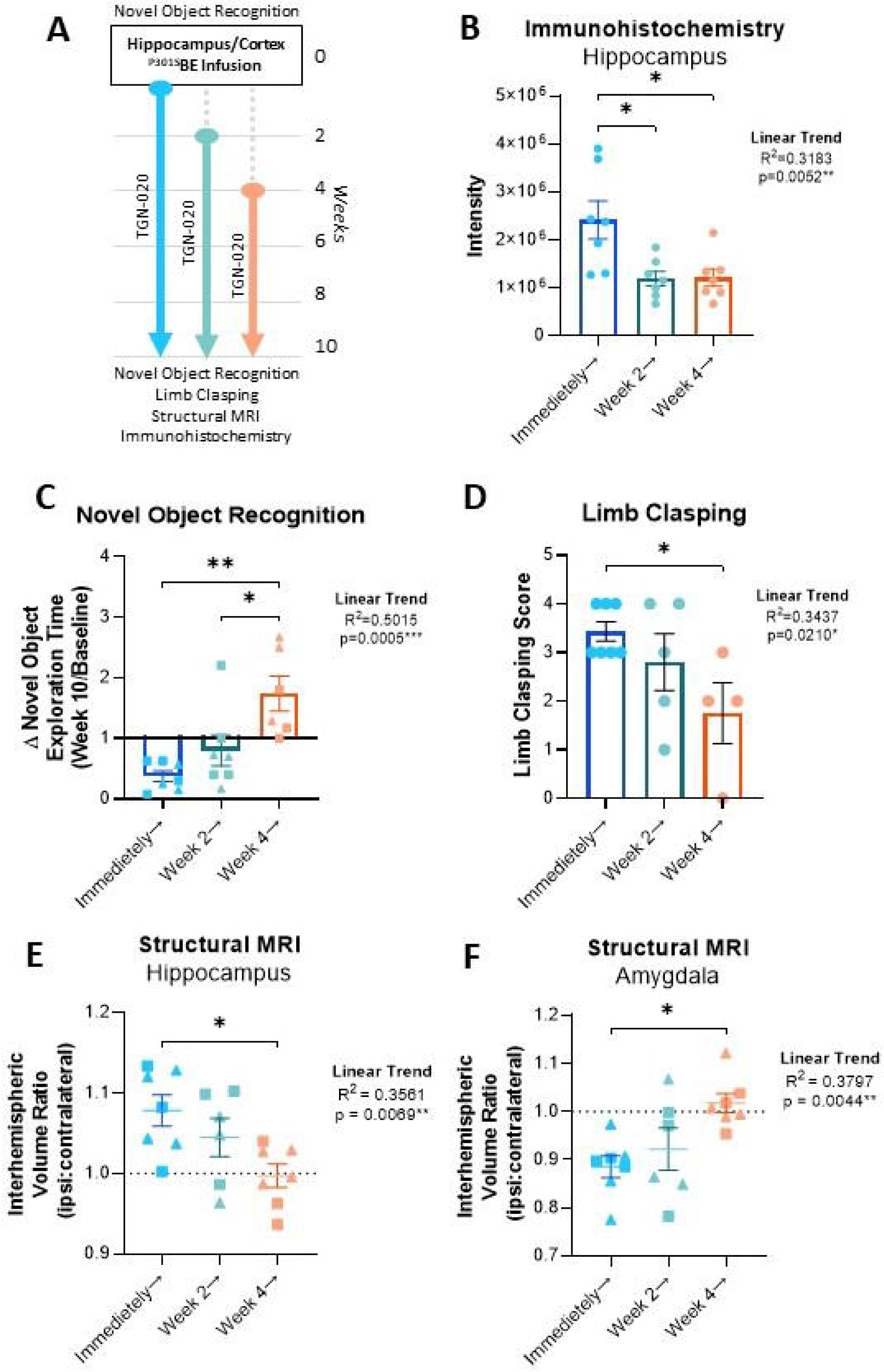
Delayed TGN-020 Treatment Impacts Its Effect on Propagating Tau Pathology. **(A)** Study design of delayed treatment experiments. Novel Object Recognition is performed prior to tau infusion, which is then followed either immediately, or 2 or 4 weeks later by chronic TGN-020 treatment. At the end of the study (week 10), the NOR task is repeated, limb clasping is assessed, and the structural MRI of the mouse brain is acquired, prior to culling for immunohistochemistry. Immunohistochemical analysis of tau (AT8) immunoreactivity in the hippocampus, reveals that immunoreactive intensity (**B**) is reduced with TGN-020 treatment delayed. This results in rescue of novel objective exploration at week 10 ( **C**) and sparing of limb clasping behaviour ( **D**) as a function of the delay in treatment. Structural MRI also reveals alleviation of volumetric differences in regions of significant ipsi>contralateral (e.g. hippocampus ( **E**)) and ipsi<contralateral (e.g. amygdala ( **F**)) difference. ▴=females, ▪=males, *=p<0.05, **=p<0.01, n=7-8 per group.

The intensity of hippocampal tau immunoreactivity in mice receiving TGN-020 immediately after tau inoculation was significantly greater than both the 2-week and 4-week delayed treatment groups (p<0.05 in both comparisons, **Fig. 5B**), suggesting a critical influence of the glymphatic system on tau burden in the initial 2-weeks following ‘seeding’. Moreover, a significant linear trend was observed between the decline in tau immunoreactivity and the delay in TGN-020 treatment, indicating clear influence of the glymphatic system on the pathological accumulation of tau in the brain.

Following 10-weeks of TGN-020 treatment, ^P301S^BE injected mice spend significantly less time exploring objects in a NOR task compared to earlier timepoints ( **Fig. 5C**, **Suppl. Fig. 7B**). However, it appeared that motivation to perform the task was spared when treatment initiation was delayed (linear trend, R^2^=0.5015, p<0.001, **Fig. 5C**); ratio of time spent exploring the novel object at week 10 compared to baseline was significantly greater in both the 2-week and 4-week delayed groups (p<0.05 and p<0.01 respectively) compared to mice which received drug treatment immediately post-inoculation. Limb clasping behaviour was also significantly alleviated by delaying drug treatment (linear trend, R^2^=0.3437, p<0.05, **Fig. 5D**) with the 4-week delay treatment group exhibiting a significantly lower limb clasping score compared to the mice treated immediately (p<0.05). Consistent with histological measures, these behavioural outcomes similarly suggest that the period immediately followed tau inoculation is pivotal for glymphatic influence on tau pathology.

Upon examination of structural MR images of mouse brains at week 10, as per our previous findings, significant tau induced interhemispheric differences were observed in the striatum and amygdala (ipsi<contra), and in the midbrain, anterior commissure, fimbria and hippocampus (ipsi>contra) (**Suppl. Fig. 8**). However, with each 2-week delay in treatment initiation, fewer brain regions appeared significantly altered between hemispheres ( **Suppl. Fig. 8**). This time dependent mitigation is particularly evident in the hippocampus ( **Fig. 5E**) and amygdala (**Fig. 5F**), illustrating the significant trends towards hemispheric equality with treatment delay in these regions (linear trends, R ^2^=0.9958 and 0.9299 respectively, p<0.01 in both).

Together these data indicate that there is a clear time window where alterations in the glymphatic system can cause a severe impact on tau propagation and pathogenesis, and that the impact of glymphatic alteration diminishes with pathological progression.

## Discussion

Growing evidence suggests a relationship between glymphatic clearance and the propagation of amyloidosis in the context of neurodegeneration, but to date a clear link between this clearance pathway in the brain and tau propagation is yet to be established. In this study, we investigated the impact of chronic glymphatic disruption on tau propagation in an animal model. Our study demonstrates that a reduction in glymphatic clearance not only leads to the advancement and dissemination of tau pathology throughout the brain, but also exacerbates atrophic changes and impairs cognitive performance. These findings strongly underscore the direct association between the glymphatic system and the propagation of tau pathology. Further, delayed glymphatic disruption experiments suggest that glymphatic clearance of propagation-prone tau in the initial stages of its propagation is vital for avoidance of tau pathogenesis. These data reinforce the idea that targeting the glymphatic system for therapeutic treatment, particularly in the early stages of disease, could delay or prevent the onset of clinical symptoms associated with neurodegenerative diseases.

We demonstrate that chronic pharmacological modulation of AQP4 efficiently influences global glymphatic function. The reliance of the glymphatic system on AQP4 channels has been extensively investigated, with many studies demonstrating that loss of AQP4 expression and polarised localisation lead to a decrease in glymphatic inflow and brain clearance (3, 7–12, 36). It is clear that AQP4 channels play a central role in the normal physiological function of the glymphatic system, however so far it has not been investigated whether chronic pharmacological blockade of these channels has a similar impact on glymphatic flow; previous studies relying on global knockout models as their main research tool. Due to the known drawbacks of using knockout models, where developmental and other compensatory variables have the potential to confound results, we used a pharmacological approach in which glymphatic function was experimentally modulated, in line with the initiation of pathology. Our experiments using TxR-d3 to evaluate glymphatic flow upon chronic treatment with a specific AQP4 inhibitor (30–32), TGN-020, show that long term blockade of these channels leads to a dramatic decrease of glymphatic flow in the brain, a phenomenon observed without the occurrence of any apparent physiological compensatory regulation of channel expression or localisation. To the best of our knowledge, this is the first time in which a study has used a pharmacological inhibitor of AQP4 to induce glymphatic downregulation chronically. This model has proven ideal to study the role of the glymphatic system in the brain free from the drawbacks of genetic knockout systems and closely resembles to the physiological state of the brain. Future use of this model system might allow evaluation of the extent of plasticity of the glymphatic system in the brain and explore whether its function and flow can be heightened or enhanced, emulating what could be a viable treatment to potentiate glymphatic clearance in disease states and with age.

We demonstrated that the extent of tau propagation is affected by glymphatic dysfunction. Tau is a microtubule associated protein crucial for neuronal biology and function, which under pathological circumstances can form toxic intracellular filamentous aggregates characteristic of several neurodegenerative diseases known as tauopathies (1, 37). Despite its intracellular location, tau is capable of transcellular propagation via release of prions into the extracellular space (2, 15). In turn, the glymphatic system is thought to be heavily involved in homeostatic regulation of the extracellular space, and management of metabolic waste clearance, implying its innate role in the clearance of extracellular tau prions. Indeed, it was recently shown using an AQP4 knockout mouse model that extracellular tau is subject to glymphatic clearance, and that glymphatic functionality protects the brain from tau aggregation and neurodegeneration (11). Likewise, we previously provided further evidence suggesting a strong relationship between glymphatic function and tau accumulation, demonstrating reduced AQP4 perivascular localisation, decreased CSF/ISF exchange and impaired glymphatic inflow and clearance of tau in a mouse model of tauopathy (14). Taken together, these data suggest a vicious cycle relationship between tau aggregation and glymphatic impairment in the brain. To further probe this relationship, here, we tested whether the glymphatic system affected not only the aggregation of tau, but its propagation in the brain. Using the P301S mouse model of tau propagation, we showed that chronic glymphatic inhibition resulted in amplified phosphorylated tau signal in the hippocampus, and intensified interhemispheric spread (i.e. propagation) in both the hippocampus and cortex. Consistent with a glymphatic mediated clearance mechanism of prion-prone extracellular tau species, this data suggests that increased extracellular tau concentration upon failure of glymphatic function is capable of aggravating tau spread throughout the brain. It has been known for some years that glymphatic failure is associated with aging (18, 19), i.e. the greatest risk factor for dementia and many other neurodegenerative diseases (38). Hence our data taken together with others’ showing that tau is more prone to spread in the aged brain, suggest that glymphatic failure with age is a likely contributor to the development and intensification of neurodegenerative disease.

Previous neuropathological characterisation of the animal model used in this study demonstrated that hippocampal/cortical tau inoculation results in spread of misfolded tau pathology, in both an anterograde and retrograde fashion, to synaptically connected brain regions (20). Consistent with these previous findings, and in line with the bilateral mode of spread of tau pathology previously seen in this model, we observed not only ipsilateral but significant contralateral tau deposition in the hippocampus and cortex, which was exacerbated upon glymphatic inhibition. In addition, our use of in depth structural MR imaging and unbiased, automated segmentation of images provides a detailed description of extra-hippocampal/cortical brain regions prone to volumetric change as a result of tau propagation in this model. Acquiring these images *in vivo*, as we have done in this study, enables detection of subtle changes in brain volumes, which are not usually accessible using post-mortem histological approaches, due to the fixation, dehydration and sectioning involved. To the best of our knowledge, the dataset which we present here provides the first brain-wide, unbiased volumetric analysis of this animal model, adding context to previously published histological datasets (e.g. 20).

In our study, we observed that inhibition of glymphatic function following hippocampal/cortical tau inoculation causes ipsilateral enlargement of connected white matter tracts, namely the anterior commissure and fimbria. These regions were observed to increase in size in the ipsilateral hemisphere (relative to the contralateral side) of tau injected groups; an effect which was intensified by drug treatment and lessened with treatment delay. The fimbria is part of the fornix, which has been shown to be increasingly burdened by ‘dot-like’ tau structures post-inoculation in this model, comparable with tau aggregates in neuronal axons (20). The ipsilateral enlargement we observed in this brain region is consistent with neuroinflammation/oedema due to tau burden, and the increased effect we see here upon drug treatment which was attenuated by treatment delay suggests that in line with other findings we report, glymphatic impairment exacerbates propagation along nerve fibre bundles. Consistently, the anterior commissures connect corresponding regions in the left and right hemisphere, and the increased effect on the volume of this region observed here with drug treatment likely reflects the amplified transhemispheric propagation we see in connected grey matter regions, e.g. the cortex; the anterior commissures acting as the conduit for transhemispheric propagation in this region. We also observe transhemispheric propagation of tau in the hippocampus, likely facilitated by the hippocampal commissures which have also been previously demonstrated to be tau positive in this model, to a similar extent to the fornix (20). It is also important to note, that in all animal groups, ipsilateral enlargement of the internal capsule was also observed; likely the physical result of intracerebral injection rather than tau itself, as similar degrees of interhemispheric inequality were observed in both ^WT^BE injected ^P301S^BE mice (**Suppl. Fig. 8**), and in ^WT^BE and ^P301S^BE injected wildtypes (data not shown), hence exclusion of this region from inter-group comparisons (i.e. **Fig. 4**).

We demonstrate here that unilateral hippocampal/cortical inoculation with tau, and subsequent glymphatic inhibition result in interhemispheric differences in regional grey matter volumes, and corresponding behavioural deficits. We show that in the hippocampus (parcellated ROI encompassing CA1, CA3, CA4, dentate gyrus, subiculum and hippocampal commissure), and the midbrain (parcellated ROI encompassing the ventral tegmental area, the substantia nigra, midbrain reticular nucleus, dorsal raphe nucleus, red nucleus, nucleus sagulum, pedunculopontine nucleus, and parabigeminal nucleus), there are significant interhemispheric differences (ipsi<contralateral) in brain volume, which are both exacerbated by drug treatment. These ipsilateral enlargements were not mirrored in either the ^WT^BE injected P301S mice, or ^P301S^BE injected wildtype mice (data not shown), and appearance of this phenomenon in ^P301S^BE injected P301S mice is therefore likely the result of active neurodegenerative processes taking place here; neuroinflammation/oedema caused by infusion of pathological tau ‘seeds’ directly/locally and resulting synaptic loss. Consistently, we have noticed in this model that astrocyte activation (GFAP expression) is ∼3.5 fold higher in the ipsilateral hippocampus over the contralateral hippocampus (data not shown), an upregulation that persists from shortly after inoculation right up until end stage (10-weeks, i.e. the timepoint at which the animals were imaged in this study). Unsurprisingly, as a result, we observed alterations in recognition memory (NOR) in these mice, which is dependent on the hippocampal formation and its cortical connections (33–35). Away from the injection site in mice, ipsilateral atrophy rather than enlargement was observed, in the amygdala and striatum parcellated ROIs, which again were exacerbated by drug treatment and attenuated as a function of treatment delay. These regions play a large role in governing motivated behaviour (39–42). Of note, the nucleus accumbens in the ventral striatum (encompassed in the ‘striatum’ parcellated ROI), has been shown to become heavily burdened in ^P301S^BE injected P301S mice (20), and is thought to play a key role in motivation (43, 44). It is not surprising then that although we did not employ tasks which directly test the function of this brain region here, in drug treated mice at week 10, there appears to be clear lack of motivation in behavioural tasks, e.g. immobility in the open field task, and lack of exploration in the NOR test, consistent with this MR finding.

Performance and motivation in behavioural tasks were less impaired when TGN-020 treatment in mice was initiated later, in line with attenuated tau accumulation and regional brain volumes. This finding in particular highlights the initial 2-weeks after tau inoculation as a critical window in which pathological development can be influenced by glymphatic alteration. This is strongly corroborative data, as extracellular tau ‘seed’ content is at its greatest immediately following intraparenchymal inoculation in this animal model, but this finding has potential implications when considering its translation to the clinic. As such, extrapolation of this data would suggest that targeting glymphatic function would be most advantageous in the early stages of disease development. Indeed, this is logical since glymphatic function has been shown to decline both with age (18, 19) and in the advanced stages of disease (13, 14, 45–50). Before this hypothesis can be tested however, much further work is required, in order to identify and fully characterise an appropriately translatable glymphatic enhancing strategy.

## Conclusions

Our data serve as crucial proof-of-concept evidence, demonstrating the ability of pharmacological modulation of glymphatic function to exert a disease-modifying effect *in vivo*. Moreover, our investigations using an animal model of ’prion-like’ tau propagation yield novel insights into the intricate relationship between glymphatic function and the underlying pathways responsible for protein propagation within the brain.

## Declarations

### Ethics approval and consent to participate

All animal work was performed in accordance with the UK’s Animals (Scientific Procedures) Act of 1986 and approved by UCL’s internal Animal Welfare and Ethical Review Board.

### Consent to publish

Not applicable.

### Availability of data and materials

The datasets used and/or analysed during the current study are available from the corresponding author on reasonable request.

### Competing interests

The authors declare that they have no competing interests.

### Funding

This work was supported by research fellowship awards from both Alzheimer’s Research UK (ARUK-RF2019A-003) and Parkinson’s UK (F-1902) made to IFH. MFL receives funding from the Rosetrees Trust and the John Black Charitable Foundation (Grant No. A2200); the Medical Research Council (Grant Nos. MR/M009092/1 and C7893/A27590); the Brain Tumour Charity (Grant No. QfC_2018_10387); the Edinburgh-UCL CRUK Brain Tumour Centre of Excellence (Grant No. C7893/A27590); the CRUK & EPSRC Comprehensive Cancer Imaging Centre at KCL and UCL (Grant Nos. C1519/A16463 and C1519/A10331).

### Authors’ contributions

DML; acquisition and analysis of data, interpretation of data, preparation of manuscript: JAW; acquisition and analysis of data: DM; analysis of data: LW; acquisition and analysis of data: DP; analysis of data: SKL; acquisition of data: ZA; conception, design of the work: MFL; conception, design of the work: IFH; conception, design of the work, acquisition and analysis of data, interpretation of data, preparation of manuscript. All authors read and approved the final manuscript.

## Acknowledgements

The authors would like to thank Prof Rohan de Silva and the late Prof Peter Davies for their kind gifts of the tau antibodies.

## Notes

### Competing Interest Statement

The authors have declared no competing interest.

