## Supplementary Material for "Glymphatic Inhibition Exacerbates Tau Propagation in an Alzheimer’s Disease Model"

**Supplementary Methods**

**Brain Extract Validation**

*ELISA*

Total tau content was quantified in brain extracts by ELISA (Human Tau (total) ELISA kit (#KHB0041, Invitrogen, UK), as per the manufacturer’s instructions. Briefly, homogenates were diluted 1:50,000 in standard diluent buffer and incubated on the capture antibody coated plate along with prepared standards (0-1 µg/ml human tau (total)) for 2 hrs. Samples were then removed and the wells were washed thoroughly prior to incubation with the biotinylated-secondary antibody for 1 hr. Wells were thoroughly washed again prior to incubation with streptavidin-HRP for 30 min. Wells were washed again and incubated with the stabilising chromogen for 30 min ahead of addition of the stop solution. Absorbance of the plate was read at 450 nm in a Varioskan LUX microplate reader (ThermoFisher Scientific, UK). Following background subtraction, the absorbance of standards was fitted to a 4^th^ order polynomial line of best fit in order calculate tau concentration in unknown samples.

*Dot Blots*

In order to confirm immunoreactivity of ^P301S^BE (and not ^WT^BE) with antibodies against common epitopes, dot blots were performed. Brain extracts were serially diluted, loaded (1.5 µl/sample) onto 0.45 µm nitrocellulose membrane and left to dry overnight at room temperature. Membranes were blocked in 5% non-fat milk in Tris-Buffered Saline with Tween-20 (0.1%) (TBST) for 3 hrs prior to incubation in primary antibody (either mouse anti-MC1 (abnormal AD tau conformation) (kind gift from the late Prof Peter Davies), mouse anti-TG5 (total tau) (kind gift from the late Prof Peter Davies), mouse anti-RD4 (1E/A6) (kind gift from Prof Rohan de Silva (1)), or mouse anti-β-Actin (ab8226, Abcam, UK), at 1:500 in 1% non-fat milk in TBST overnight at 4 °C. Membranes were then washed and incubated in goat anti-mouse HRP-conjugated secondary antibody (ab6789, Abcam, UK), 1:1000 in 1% non-fat milk in TBST for 3 hrs at room temperature. Membranes were washed and then incubated with chemiluminescent substrate (SuperSignal™ West Pico PLUS, ThermoFisher Scientific, UK) and imaged on an ImageQuant 800 multimode gel imager (Cytiva (GE Healthcare), UK).

*Thioflavin T Incubation*

The presence of amyloid structures in ^P301S^BE (and not ^WT^BE) was confirmed by fluorescence upon incubation with Thioflavin T. For this, brain extracts were diluted to a final concentration of 1:100 in 12.5 µM Thioflavin T in PBS. Solutions were plated in triplicate in a black, clear bottomed 96-well plate and fluorescence measured in wells on a Varioskan LUX microplate reader (ThermoFisher Scientific, UK), with excitation at 444 nm and emission at 478 nm. Background (12.5 µM Thioflavin T in PBS) readings were subtracted, and signal intensities (a.u.) directly reported.

**AQP4 mRNA Expression Analysis**

AQP4 expression was measured in the ipsilateral hippocampus of study mice at various stages throughout the pathological progression of tau, in order to confirm its continued expression with disease development. Mice received intracerebral infusion with either ^WT^BE or ^P301S^BE (see main text materials and methods) and were left to age for the appropriate time period, after which animals were killed by overdose with sodium pentobarbital (10 ml/kg, given intraperitoneally), the brain was removed from the skull and the ipsilateral hippocampus rapidly dissected out. Excised tissue was snap frozen immediately on dry ice and stored at -80 °C until further processing. mRNA was extracted from brain samples using the PureLink™ RNA Mini Kit (ThermoFisher Scientific, UK) as per the manufacturer’s instructions, which yielded RNA with a 260/280 absorbance ratio of 2.04 (±0.04). RNA was converted to cDNA using the QuantiTect Reverse Transcription Kit (Qiagen, UK) as per the manufacturer’s instructions, which was then quantified by qRT-PCR using TaqMan™ Universal PCR Master Mix and TaqMan™ probes against AQP4 (Mm00802131_m1), using GAPDH (Mm99999915_g1) and ACTB (Mm02619580_g1) as housekeeper controls (all ThermoFisher Scientific, UK), using the 2^-ΔΔCt^ method for analysis (2), normalising to wildtype baseline data.

**Brain and CSF Extraction and Analysis**

For drug efficacy study of TGN-020 in relation to brain tau clearance, after drug treatment, CSF was extracted from mice for quantification of tau content following its intracerebral infusion. For this, mice were treated with TGN-020 (either 25, 50, 100 or 200 mg/kg in 20mg/kg saline or 20mg/ml saline alone, i.p.) ahead of being anesthetized with 2% isoflurane in O_2_ at a delivery rate of 1 l/min and positioned in a stereotaxic frame. A midline incision was made on the top of the head to expose the skull and a burr hole made with a microdrill above the injection location (+2.5 mm anteroposterior, and +2 mm mediolateral to bregma). 2.5 µl of either ^P301S^BE or artificial CSF (control) was infused at a rate of 0.5 µl/min into the hippocampus (-1.8 mm ventrodorsal to the brain surface respectively) using a 5 µl Hamilton syringe, 15 min after drug treatment. A midline incision was then made at a midpoint between the skull base and the occipital margin to the first vertebrae. The underlying muscles were parted to expose the atlanto-occipital membrane and dura mater overlaying the cisterna magna, which was cleaned with saline. A durotomy was performed 15 min after intracerebral injection, using a 23-gauge needle, allowing free-flowing CSF to be collected using a narrow bore pipette tip. At the end of this procedure, mice were killed by overdose with sodium pentobarbital (10 ml/kg, given intraperitoneally). The brain was removed from the skull and snap frozen immediately on dry ice and stored at -80 °C until further processing. The collected CSF was centrifuged briefly to pellet any red blood cell contamination. Spectrophotometric analysis (417 nm, NanoDrop ND-1000, Fisher Scientific, UK) of hypotonically disrupted blood cell contaminants (freeze-thawing of rehydrated blood pellets) measured blood contamination to be <0.02% in all samples. Total tau concentration in CSF samples and homogenised (10% (w/v) in sterile phosphate-buffered saline (containing protease inhibitor cocktail, phosphatase inhibitor cocktails I and II (Sigma, UK), at a final dilution of 1:100, and 1 mM phenylmethylsulphonyl fluoride)) brain tissue was quantified by ELISA (Human Tau (total) ELISA kit (#KHB0041, Invitrogen, UK)) as above. CSF tau concentration (ng/ml), and total brain tau (ng), are reported.

**Supplementary Figures**

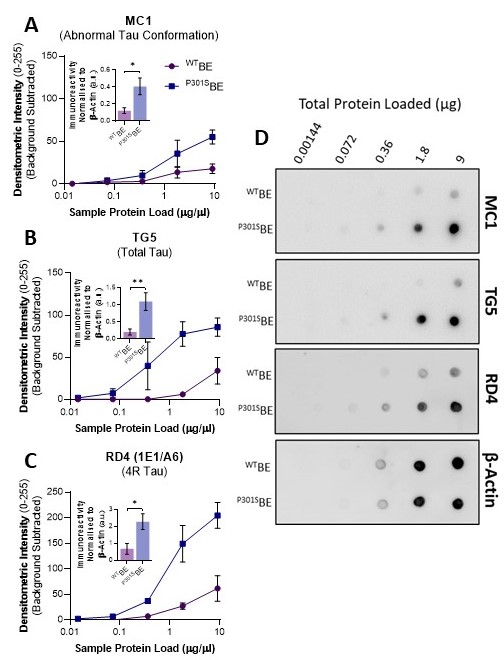

**Supplementary Figure 1 – Dot Blot Immunoreactivity Validation of Brain Extracts**

Densitometric intensity analysis of brain extract dot blots illustrating preferential immunoreactivity of ^P301S^BE over ^WT^BE with (**A**) MC1, (**B**) TG5, and (**C**) RD4 antibodies. Graphical insets illustrate differences between immunoreactivity of ^P301S^BE and ^WT^BE when signals are normalised to β-Actin across loading amounts. (**D**) Representative example dot blots used to produce data shown in (**A**-**C**). *=p<0.05, **=p<0.01.

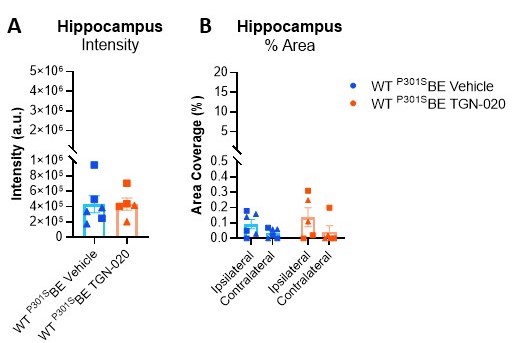

**Supplementary Figure 2 – ^P301S^BE Injection Does Not Result in Tau Pathology in Wildtype Mice**

Hippocampal AT8 immunopositive intensity (**A**) and % area coverage (**B**) in wildtype mice injected with ^P301S^BE and treated with vehicle or TGN-020, illustrating the lack of significant tau pathology induced by ^P301S^BE infusion in the wildtype mouse brain. Y-axes are included at the same scale as the corresponding data from transgenic mice (Fig. 2B, C), illustrating the difference in tau level detected. ▲=females, ■=males. n=4-5 per group.

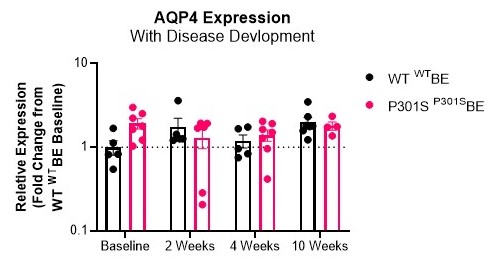

**Supplementary Figure 3 – AQP4 Expression with the Onset of Tau Propagation**

Hippocampal AQP4 (mRNA) expression measurements (qRT-PCR) indicate that AQP4 expression remains relatively constant between genotypes, and throughout pathological development in the ^P301S^BE injected transgenic mice.

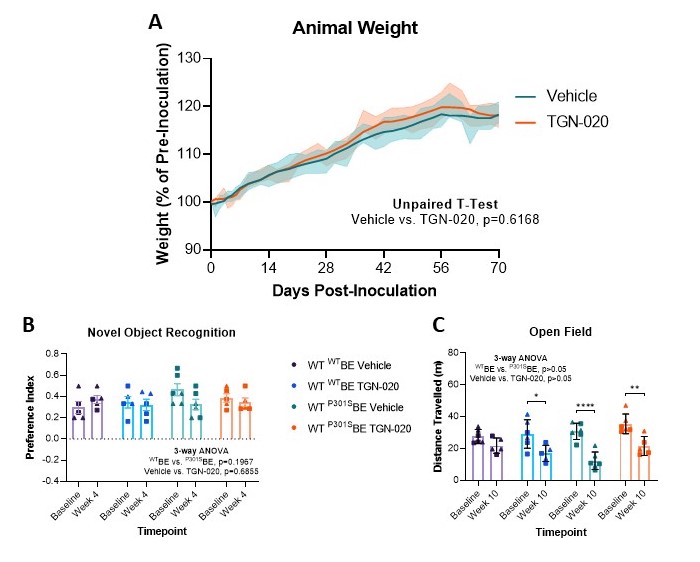

**Supplementary Figure 4 – No Effect of Chronic TGN-020 Treatment in Control (Wildtype) Mice**

(**A**) Consistent with animal growth, both vehicle and TGN-020 treatment groups (n=10-11) increase in weight over the treatment period (70 days starting from 2-months-of-age). No effect on animal weight was observed with drug treatment. (**B**) Neither tau inoculation nor TGN-020 treatment affected wildtype animal performance in the novel object recognition task (n=5-6). (**C**) Likewise, aside from the reduced motility observed by animals at week 10 compared to their baseline performance, due to test arena familiarisation, no effect was observed as a result of tau inoculation or TGN-020 treatment in mice (n=5-6). ▲=females, ■=males. *=p<0.05, **=p<0.01, ****=p<0.0001.

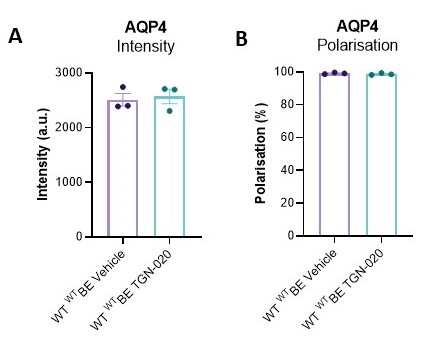

**Supplementary Figure 5 – AQP4 Expression and Polarisation Are Unaffected Following Chronic TGN-020 Treatment**

Expression measurements of hippocampal AQP4 protein (**A**) and its polarised location (**B**) (immunohistochemistry) are unaffected by chronic (10 weeks) TGN-020 treatment.

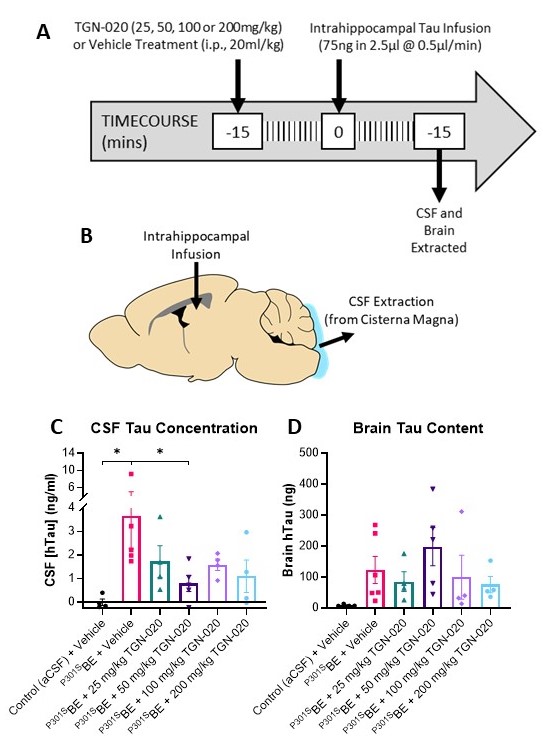

**Supplementary Figure 6 – Reduction of Brain to CSF Tau Clearance Following TGN-020 Treatment**

(**A**) Experimental timeline and (**B**) schematic of brain injections/extractions. Mice were pre-treated (15 min prior) with either TGN-020 (at varying doses) or vehicle, ahead of hippocampal infusion of tau (^P301S^BE). 15 min later, CSF samples and brains were extracted, and their tau content quantified by ELISA; CSF and brain tau content following various TGN-020 treatments is shown in (**C**) and (**D**) respectively. As expected, significantly greater tau levels were detected in CSF extracted from ^P301S^BE (vehicle) injected mice compared to aCSF (control) injected mice. This CSF tau content was reduced in all TGN-020 treated animal groups, reaching significance compared to vehicle treated mice at 50 mg/kg TGN-020. A similar, inverse trend was observed in brain tau data, with 50 mg/kg TGN-020 treated mice exhibiting a trend towards an increase in brain tau content. Based on this data, 50 mg/kg TGN-020 was taken forward for chronic treatment studies. *=p<0.05, n=4-6 per group.

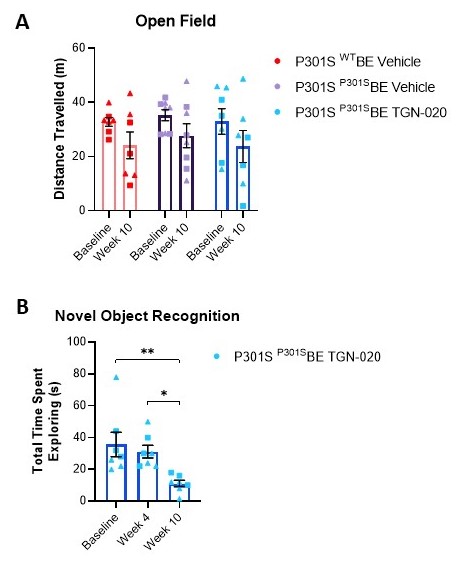

**Supplementary Figure 7 – Chronic TGN-020 Treatment Leads to Motor Dysfunction in Tau Seeded Mice**

No inter- or intra-group differences were observed in distance travelled in an open field arena at baseline and after 10 weeks of treatment (**A**). (**B**) Time spent by P301S ^P301S^BE injected TGN-020 treated mice exploring objects in the novel object recognition task considerably declined at week 10, resulting in exclusion of this timepoint from analysis. ▲=females, ■=males, *=p<0.05, **=p<0.01, n=5-8 per group.

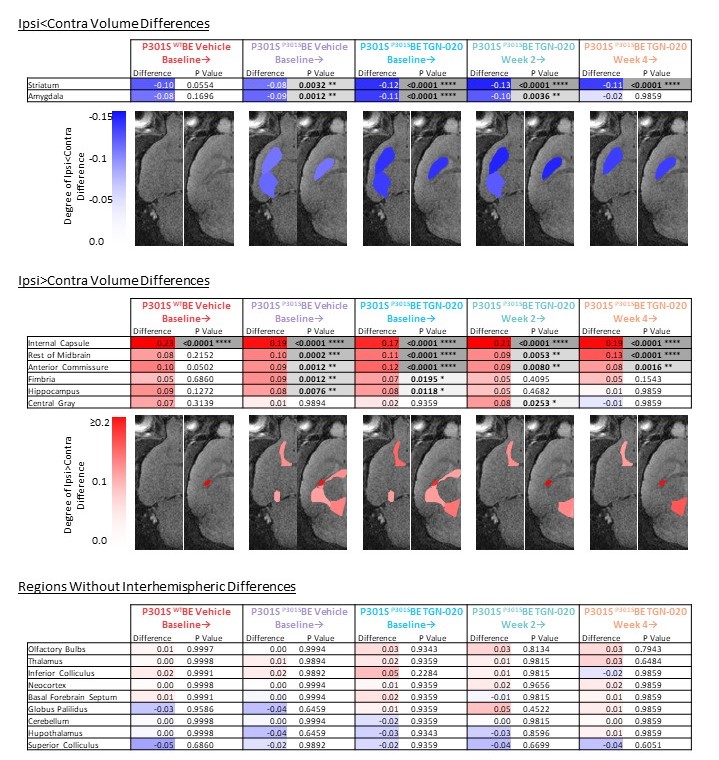

**Supplementary Figure 8 – Groupwise Interhemispheric Volume Ratio Differences**

For each brain region in each group, the interhemispheric volume ratio (ipsilateral/contralateral volume) was calculated and compared to hemispheric equality (interhemispheric volume ratio of 1) in order to appreciate the magnitude of groupwise interhemispheric inequality. Difference cells are colour coded based on the degree of difference: red denoting ipsilateral>contralateral volume, blue denoting contralateral>ipsilateral volume. Cells containing the resulting p values (adjusted for multiple comparisons) from multiple t-tests are also colour coded (greyscale) based on the level of statistical significance. For additional visualisation of data, significant regions (either ipsilateral>contralateral (red) or contralateral>ipsilateral (blue)) are displayed on a single hemisphere of horizontal MR images at two levels (left, -4 from Bregma; right, 2mm from Bregma), with colour bars denoting the degree of interhemispheric difference.
